# Stimulus degradation impairs performance in a rodent continuous performance test

**DOI:** 10.1101/2021.08.31.458320

**Authors:** Adrienne C. DeBrosse, Ye Li, Robyn Wiseman, Racine Ross, Sy’Keria Garrison, Henry L. Hallock, James C. Barrow, Keri Martinowich, Gregory V. Carr

## Abstract

Sustained attention is a core cognitive domain that is often disrupted in neuropsychiatric disorders. Continuous performance tests (CPTs) are the most common clinical assay of sustained attention. In CPTs, participants produce a behavioral response to target stimuli and refrain from responding to non-target stimuli. Performance in CPTs is measured as the ability to discriminate between targets and non-targets. Rodent versions of CPTs (rCPT) have been developed and validated with both anatomical and pharmacological studies, providing a translational platform for understanding the neurobiology of sustained attention. In human studies, using degraded stimuli (decreased contrast) in CPTs impairs performance and patients with schizophrenia experience a larger decrease in performance compared to healthy controls. In this study, we tested multiple levels of stimulus degradation in a touchscreen version of the CPT in mice. We found that stimulus degradation significantly decreased performance in both males and females. The changes in performance consisted of a decrease in stimulus discrimination, measured as d’, and increases in hit reaction time and reaction time variability. These findings are in line with the effects of stimulus degradation in human studies. Overall, female mice demonstrated a more liberal response strategy than males, but response strategy was not affected by stimulus degradation. These data extend the utility of the mouse CPT by demonstrating that stimulus degradation produces equivalent behavioral responses in mice and humans. Therefore, the degraded stimuli rCPT has high translational value as a preclinical assay of sustained attention.

## 1. Introduction

Sustained attention, the ability to focus on tasks over extended periods of time, is a fundamental cognitive domain and it is impaired in many neuropsychiatric disorders (Huntley, Hampshire, Bor, Owen, & Howard, 2017; J. Liu et al., 2021; S. K. Liu et al., 2002). In schizophrenia specifically, when controlling for the severity of other symptoms, functional outcome is correlated with sustained attention function (Green, Kern, Braff, & Mintz, 2000). Therapeutics that improve sustained attention have been approved for treating attention-deficit/hyperactivity disorder (ADHD), but the most effective of these therapies, amphetamine and methylphenidate, have substantial abuse liability and are contraindicated in other disorders characterized by attention deficits, including schizophrenia (Berman, Kuczenski, McCracken, & London, 2009). Therefore, novel therapeutics for attention deficits are needed.

Continuous performance tests (CPTs), first developed to identify attention deficits in patients with brain lesions (Beck, Bransome, Mirsky, Rosvold, & Sarason, 1956), are the most commonly used clinical measures of sustained attention (Nuechterlein et al., 2015; Riccio, Reynolds, Lowe, & Moore, 2002). CPTs are characterized by target and non-target stimuli where participants are asked to respond to targets and refrain from responding to non-targets. Trials are separated by short inter-trial intervals during which no stimuli are present. Performance is usually analyzed using components of signal detection theory, specifically the composite measure of sensitivity (stimulus discrimination) called d’ (Green and Swets, 1966). CPTs are sensitive to the attentional deficits present across multiple neuropsychiatric disorders including ADHD, schizophrenia, and major depressive disorder (Berger, Slobodin, & Cassuto, 2017; Koetsier et al., 2002; Nuechterlein et al., 2015).

Different versions of the CPT have been designed to test particular components of sustained attention (Borgaro et al., 2003). Degraded stimuli versions, where the contrast between the stimuli and background is decreased, have been used to test the stimulus detection and discrimination components of information processing in the CPT (Nuechterlein, Parasuraman, & Jiang, 1983). In general, stimulus degradation impairs performance in the CPT by making stimulus discrimination more difficult and potentiating time on task decrements in vigilance (Grier et al., 2003). Patients with schizophrenia tend to be more sensitive to stimulus degradation effects than other participants (Nuechterlein, Edell, Norris, & Dawson, 1986; Nuechterlein et al., 2015).

Recently, a mouse version of the CPT using touchscreen-based operant chambers was developed and validated (Kim et al., 2015). In the ensuing years, this CPT has been used to test the procognitive effects of psychostimulants and identify brain regions involved in sustained attention in mice (Caballero-Puntiverio, Lerdrup, Arvastson, Aznar, & Andreasen, 2020; Caballero-Puntiverio et al., 2019; Hvoslef-Eide et al., 2018). Kim and colleagues showed that decreasing image contrast in the touchscreen CPT impairs performance (Kim et al., 2015). Here, we extend that result by including an increased number of degradation levels and demonstrate the translational utility of this CPT procedure by showing that mice performance is altered in similar ways compared to human performance in the DS-CPT across multiple behavioral measures, including overall sensitivity and reaction time variability.

## 2. Materials and methods

### 2.1. Mice

Eight-week-old male C57BL/6J mice (Strain #: 000664; The Jackson Laboratory, Bar Harbor, ME, USA) were used in all experiments. The mice were group-housed in disposable polycarbonate caging (Innovive, San Diego, CA, USA) and maintained on a 12/12 light/dark cycle (lights on at 0600 hours). Water was available in the home cage *ad libitum* throughout all experiments. The mice were fed Teklad Irradiated Global 16% Protein Rodent Diet (#2916; Envigo, Indianapolis, IN, USA) in the home cage *ad libitum* until the start of the food restriction protocol. Two separate cohorts of mice (Cohort A, n = 7/male and 7/female; Cohort B, n = 7/male and 7/female) were tested in these experiments and all testing was done Monday-Friday during the light phase (1200-1600 hours). All experiments and procedures were approved by the Johns Hopkins Animal Care and Use Committee and in accordance with the *Guide for the Care and Use of Laboratory Animals*

### 2.2 Food Restriction Protocol

Upon arrival at the animal facility, mice were given at least 72 hours to acclimate to the colony room before handling by experimenters. Mice were handled and weighed daily from that point forward. After at least two days of handling, mice were food restricted to 3g of chow per mouse per day to maintain 85-90% of their predicted free-feeding weight based on average growth-curve data for the strain (The Jackson Laboratory). To familiarize the mice with the Nesquik ^®^ strawberry milk (Nestlé, Vevey, Switzerland) reward used in the rCPT, we introduced the milk to the home cage on 4×4 inch weighing paper (VWR, Radnor, PA, USA). The weighing paper was left in the cage until all mice had sampled the strawberry milk. This procedure was repeated for a total of two days.

### 2.3 rCPT procedure

#### 2.3.1 Apparatus

Eight Bussey-Saksida mouse touchscreen chambers (Lafayette Instruments, Lafayette, IN, USA) running ABET II software (Campden Instruments, Loughborough, UK) were used for the behavioral testing.

#### 2.3.2 rCPT Training

##### 2.3.2.1 Habituation

Mice were given 30-min habituation sessions to acclimate them to the touchscreen chambers. In habituation sessions, 1 mL of strawberry milk was placed into the reward tray. The screen was responsive to touch, but touches were not rewarded. Mice were advanced to the next stage of training following three habituation sessions with at least one session where the mouse had consumed all of the strawberry milk.

##### 2.3.2.2 Stage 1

In Stage 1, mice were trained to touch a white square. Each session lasted for 45 minutes. The square was displayed for 10 seconds at a time, defined as the stimulus duration (SD) followed by a 0.5 second limited hold (LH) period during which the screen was blank, but a touch would still yield a reward. Upon interacting with the stimulus, a one-second tone (3 kHz) would sound, a small amount of reward would be dispensed, and the reward tray would be illuminated. Head entry into the reward tray was detected by an IR beam, and following head entry the 2-sec intertrial interval (ITI) would begin. If the mouse did not interact with the stimulus during the SD or LH, the ITI would start and the next trial would follow. The criterion for advancement to Stage 2 was for a mouse to obtain 60 rewards within a single session.

##### 2.3.2.3 Stage 2

In Stage 2, the white square pattern was replaced with either horizontal or vertical bars. Sessions were still 45 minutes long. Each mouse was assigned one of the stimuli and this would be that mouse’s target, or S+, for the duration of the experiment and this S+ assignment was counterbalanced for each group. The SD was reduced from 10 seconds to 2 seconds, and LH was increased from 0.5 seconds to 2.5 seconds. As in Stage 1, criterion for advancement was 60 rewards earned within a single session.

##### 2.3.2.4 Stage 3

In Stage 3, a non-target (S-) was added. On each trial, there was a 50% chance of either S+ or S-presentation. SD and LH were identical to stage 2, but the ITI was increased to 5 seconds. Screen touches during S-trials would not yield a reward and would start the ITI. The Stage 3 criteria for advancement were a minimum of seven sessions, during which at least two consecutive sessions had a d’ score of 0.6 or higher. The d’ discrimination index and other performance metrics are described in Table 1. Mice that reached criterion on days other than Friday were held on Stage 3 training until the following Monday.

| Parameter | Symbol | Formula | Description |
| --- | --- | --- | --- |
| Hit rate | HR | Hits/(Hits + Misses) | Correct response to S+ |
| False alarm rate | FAR | False alarms/(False alarms + Correct rejection) | Incorrect response to S- |
| Sensitivity | d' | $z(HR) - z(FAR)$ | Measure of ability to discriminate between S+'s and S-'s |
| Response criterion | c | $-(z(HR) + z(FAR))/2$ | Measure of response bias; higher values corresponds to more conservative biases and vice versa |

$z$ = the inverse of the cumulative distribution function for the Gaussian distribution

##### 2.3.2.5 Degraded Stimuli Testing

Following completion of Stage 3, mice in Cohort A moved to the degraded stimuli test phase. Mice were exposed to four levels of degraded stimuli (50, 75, 87.5, and 93% degraded; Figure 1). Degradation was achieved by editing the images using the add noise filter in Photoshop (Version 2020, Adobe, San Jose, CA, USA). We randomly changed a percentage of the pixels in each image according to a Gaussian distribution. Degraded stimuli were the S+ and S-images from Stage 3 where the degradation percentage represents the percentage of pixels replaced (Figure 1). Degraded stimuli sessions were identical to Stage 3 except for the S+ and S-. Mice completed baseline Stage 3 sessions on Monday, Wednesday, and Friday and degraded stimuli sessions on Tuesday and Thursday. The degradation level order was counterbalanced within each group.

**Figure 1.**
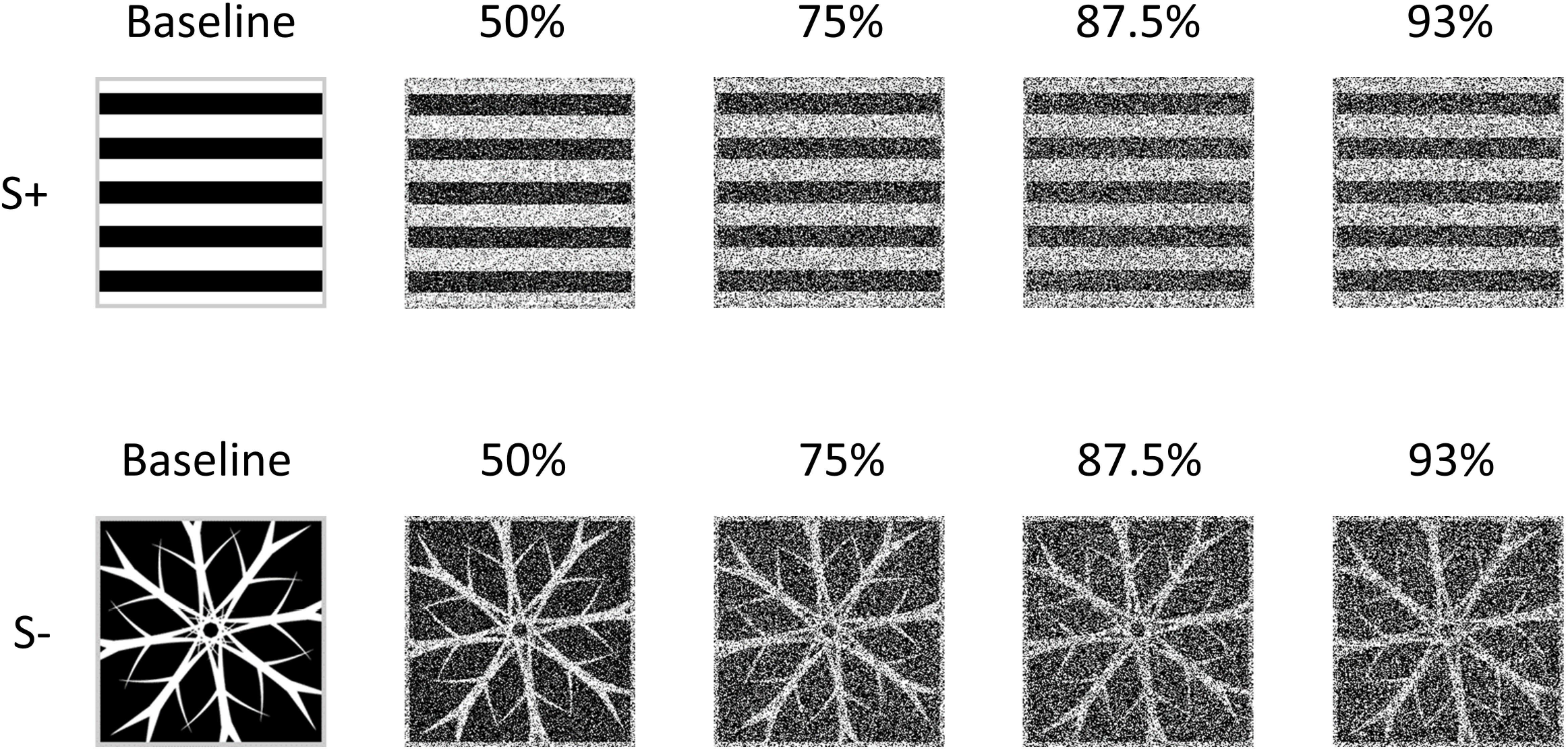
Stimulus exemplars. S+’s consisted of either horizontal (shown) or vertical (not shown) arrays of black and white boxes. The S-was a snowflake pattern. Stimulus degradation was achieved by randomizing a set percentage of the pixels in the image according to a Gaussian distribution.

### 2.4 Data analysis

Experiment databases were pulled from ABET II (Lafayette Instruments, Lafayette, IN, USA) and initial processing was done in Excel. This initial processing involved averaging the two baseline, non-degraded stimulus sessions from the day before and after each degraded stimulus session. Prism 9 (GraphPad Software, LLC, San Diego, CA, USA) was used for all data analysis. T-tests and ANOVAs were used for analysis where appropriate. The significance threshold was set at *p* < 0.05. Sidak post hoc tests were used where appropriate.

## 3. Results

### 3.1 No sex difference in sessions to criterion during training

Table 2 shows the summary data for the DS-CPT training stages. We found no difference between males and females on the number of sessions to reach criterion in Stage 1 (*t*_*26*_ = 0.6662, *p* = 0.5112), Stage 2 (*t*_*26*_ = 1.367, *p* = 0.1834), or Stage 3 (*t*_*26*_ = 0.3707, *p* = 0.7139). Because there were two conditions for completion of Stage 3, we also analyzed the number of sessions to reach the criterion of two consecutive sessions with a d’ value above 0.6. Here, we also found no difference between males and females on the Stage 3 d’ criterion (*t*_*26*_ = 1.436, *p* = 0.1629).

**Sessions to criterion**
|  | Stage 1 | Stage 2 | Stage 3 | Stage 3 d' |
| --- | --- | --- | --- | --- |
| Male | 5.79 ± 1.13 | 1.07 ± 0.07 | 7.64 ± 0.64 | 4.93 ± 0.83 |
| Female | 4.93 ± 0.62 | 1.57 ± 0.36 | 7.93 ± 0.43 | 6.36 ± 0.55 |

### 3.2 Male and female mice have similar baseline performance, but different response strategies

Although female mice advanced through training at a similar pace compared to male mice, we noticed that female mice trended toward lower d’ values during early training sessions. We analyzed the last five Stage 3 sessions for a formal comparison. We chose to focus on the last five sessions because mice did not complete the same number of sessions, but all mice had d’ values above 0.6 on their final session. We found that performance improved across the five sessions for both males and females (*F*_*4, 104*_ = 29.87, *p* < 0.0001; Figure 2a) and that females had lower d’ values across the sessions (*F*_*1, 26*_ = 5.051, *p* = 0.0333). There were no session X sex interactions (*F*_*4, 104*_ = 1.250, *p* = 0.2944). Over these five sessions, female mice showed a more aggressive response strategy quantified by a lower response criterion, c (*F*_*1, 26*_ = 4.803, *p* = 0.0376; Figure 2b). Response criterion was stable over time as there were no effects of session (*F*_*4, 104*_ = 0.9266, *p* = 0.4515) or session X sex interactions ((*F*_*4, 104*_ = 2.143, *p* = 0.0807). Despite the lower d’ scores for females early in training, males and females had similar scores by the end of Stage 3 (Figure 2c). We compared the performance of male and female mice on their final Stage 3 session and found no significant effect of sex on performance (*t*_*26*_ = 0.8522, *p* = 0.4019). Female mice still exhibited a more liberal response profile, quantified as a lower c value (Figure 2d, *t*_*26*_ = 2.240, *p* = 0.0338). The more liberal response strategy in females is not due to a difference in hit rates between males and females (Figure 2e, *t*_*26*_ = 1.424, *p* = 0.1633), but results from a significantly higher false alarm rate in females (Figure 2f, *t*_*26*_ = 3.555, *p* = 0.0015).

**Figure 2.**
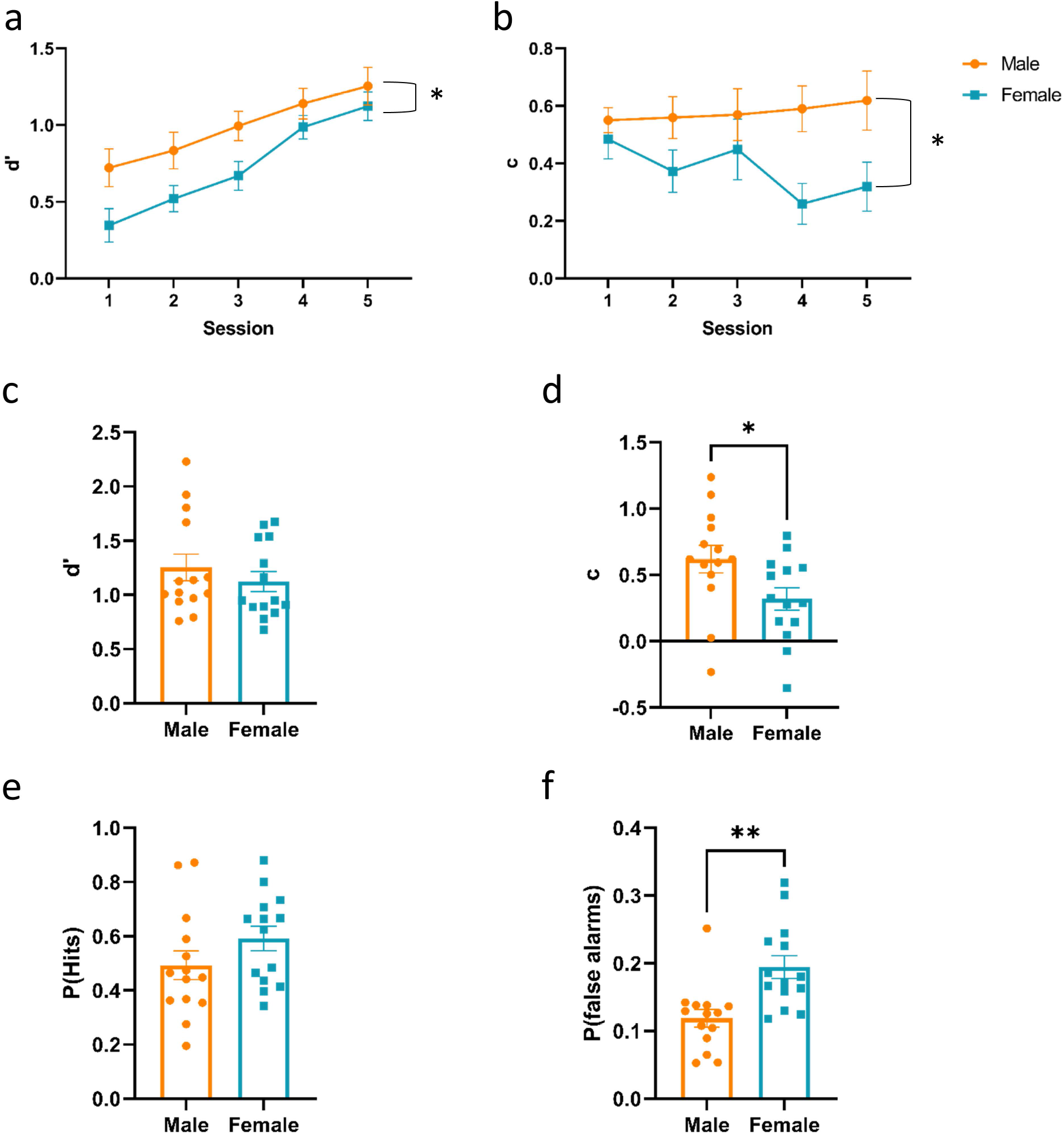
CPT performance during Stage 3 training. (a) Discrimination (d’) improved across the last five sessions of training and males produced higher d’ values compared to females across the five sessions. (b) Females display a more liberal response criterion compared to males. In the final training session, there was no difference on discrimination between male and female mice (c), however, the difference in response criterion was still present (d). The difference in response criterion was not driven by a difference in the hit rate (e), but it appears to be driven by a significant difference in the false alarm rate (f). n = 14/group and data are represented as the mean ± SEM. * *p* < 0.05; ** *p* < 0.01.

### 3.3 Degraded stimuli impair CPT performance

We next tested the effects of degraded stimuli on performance in Cohort A. We found degraded stimuli significantly impaired performance as measured by d’ in both male and female mice (*F*_*4, 48*_ = 12.10, *p* < 0.0001; Figure 3a). There was no effect of sex on performance (*F*_*1, 12*_ = 2.290, *p* = 0.1561) or any degradation X sex interactions (*F*_*4, 48*_ = 0.4244, *p* = 0.7903). Post hoc analyses showed that each of the degradation levels (50, 75, 87.5, and 93%) significantly impaired performance compared to baseline.

**Figure 3.**
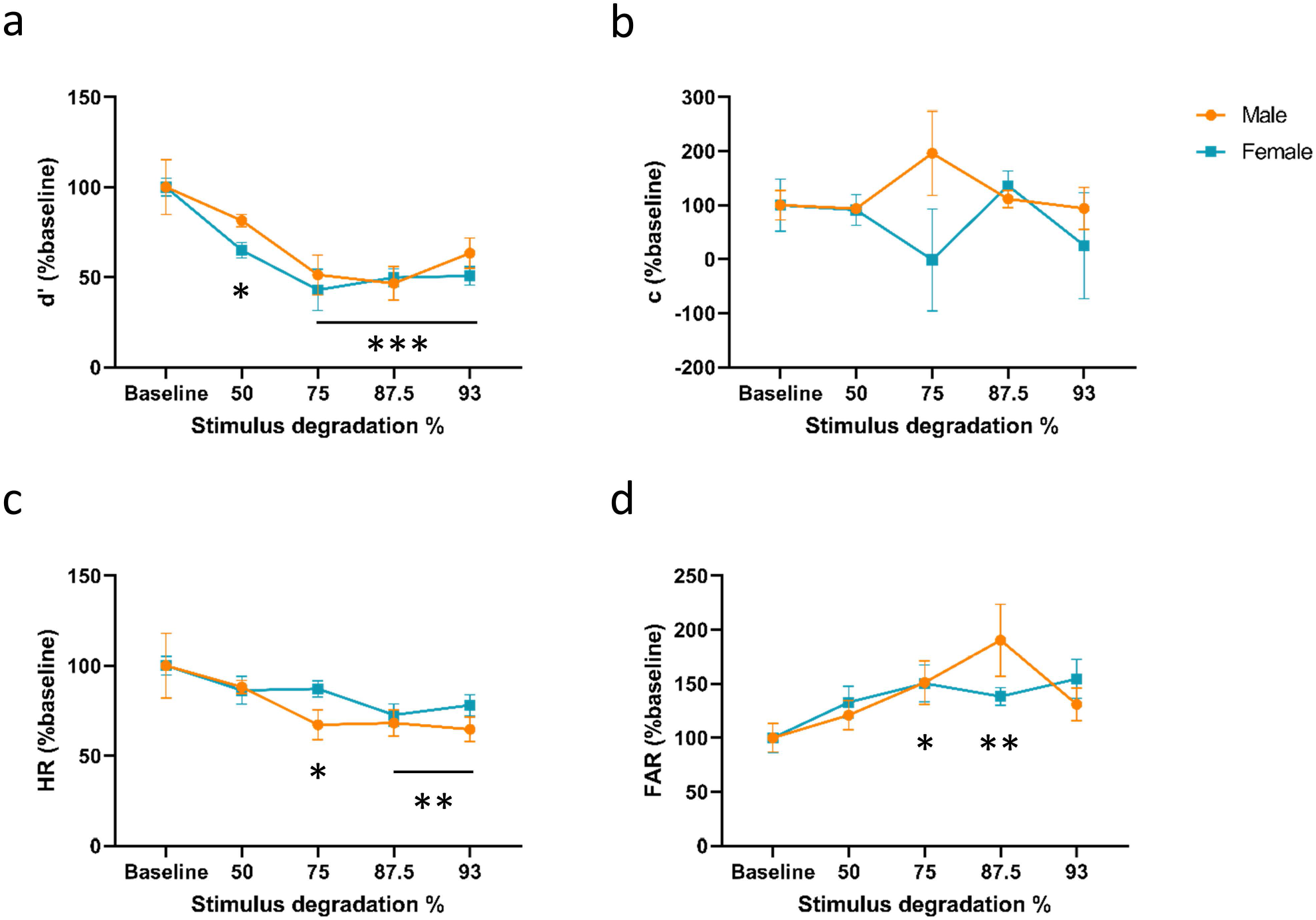
CPT performance during stimulus degradation testing. (a) d’ is significantly decreased by stimulus degradation at all levels tested. (b) There was no effect of stimulus degradation on response criterion, but degradation did decrease the hit rate (c) and increased the false alarm rate (d). n = 7/group and data are represented as the mean± SEM. \**p* < 0.05; \*\**p* < 0.01; \*\*\**p* < 0.001 compared to the baseline condition.

During training, female mice demonstrated a more liberal response strategy compared to male mice. The liberal response strategy was characterized by an increased false alarm rate and no change in the hit rate. Here, we normalized the response criterion value within sex and analyzed the effect of stimulus degradation on this measure. Degraded stimuli had no effect on response strategy in either males or females (*F*_*4, 48*_ = 0.3225, *p* = 0.8615; Figure 3b). There were also no degradation X sex interactions (*F*_*4, 48*_ = 1.232, *p* = 0.3098). As expected with a decrease in d’ and no change in c, degradation significantly decreased the hit rate (*F*_*4, 48*_ = 5.851, *p* = 0.0006; Figure 2d) and increased the false alarm rate (*F*_*4, 48*_ = 4.123, *p* = 0.0060; Figure 3c).

### 3.4 Stimulus degradation increases response reaction times and reaction time variability

In human versions of the CPT, degraded stimuli are associated with longer reaction times. Similarly, with our mouse version, we found response times on hit trials increased as degradation level increased (*F*_*4, 48*_ = 3.605, *p* = 0.0120; Figure 4a). Female mice had faster reaction times across all degradation levels (*F*_*1, 12*_ = 7.988, *p* = 0.0153) and there were no significant degradation X sex interactions (*F*_*4, 48*_ = 0.4259, *p* = 0.7892).

**Figure 4.**
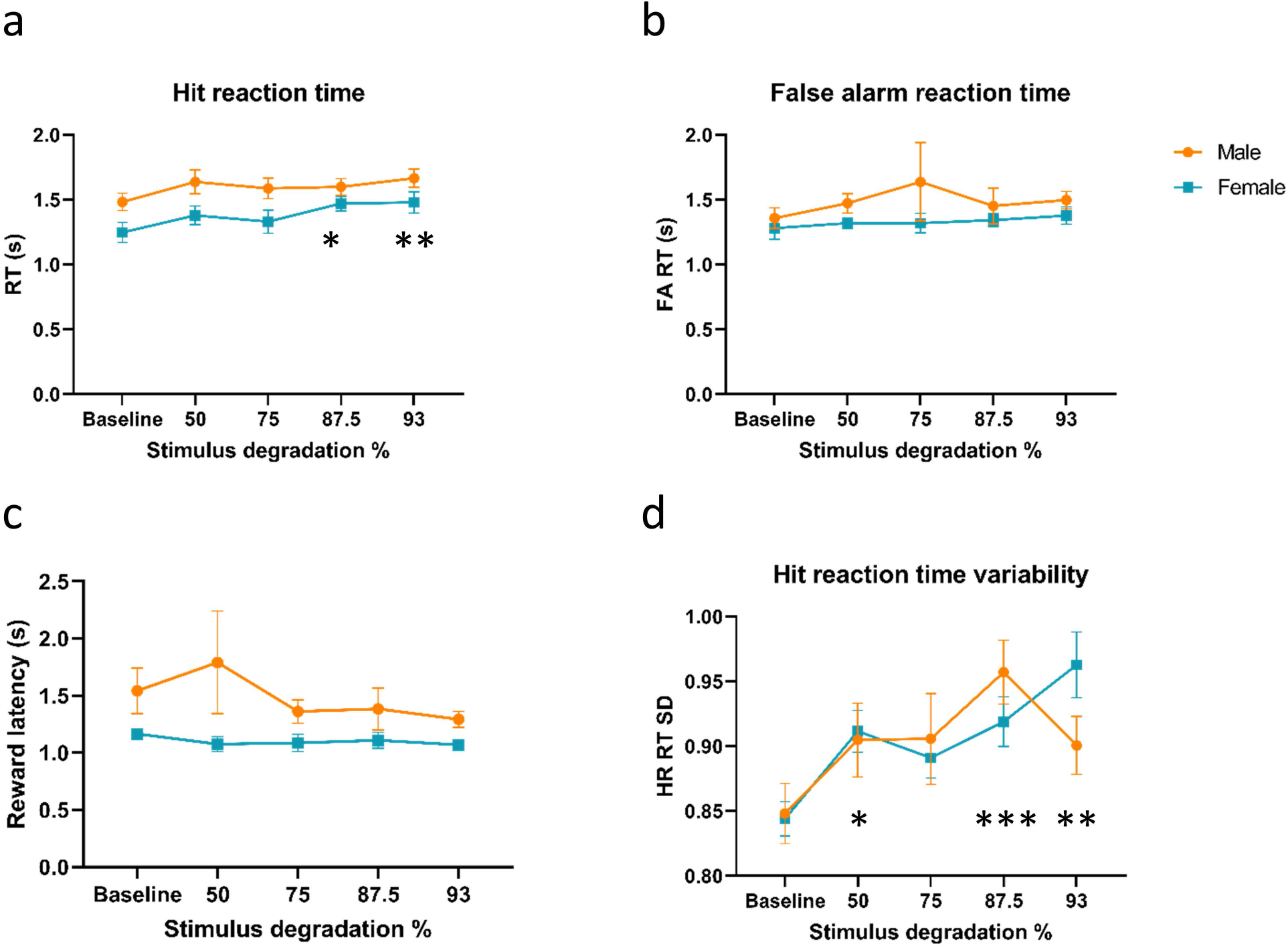
Reaction times during stimulus degradation testing. (a) Hit reaction time increases as stimulus degradation increases while false alarm reaction time (b) and reward latency (c) are unaffected. Additionally, the hit reaction time variability (d), measured as the SD of the hit reaction time increases with stimulus degradation. n = 7/group and data are represented as the mean ± SEM. \**p* < 0.05; \*\**p* < 0.01; \*\*\**p* <0.001 compared to the baseline condition.

We next tested whether the effects of degradation and sex on reaction times during hit trials carried over to false alarm trials and latencies to collect reward following correct trials. There were no significant effects of degradation (*F*_*4, 48*_ = 0.6248, *p* = 0.6471), sex (*F*_*1, 12*_ = 2.042, *p* = 0.1786), or degradation X sex interactions (*F*_*4, 48*_ = 0.4097, *p* = 0.8008; Figure 4b) on false alarm trials. Moreover, there were no effects of degradation (*F*_*4, 48*_ = 1.346, *p* = 0.2668), sex (*F*_*1, 12*_ = 3.839, *p* = 0.0737), or degradation X sex interactions (*F*_*4, 48*_ = 1.247, *p* = 0.3038; Figure 4c) on reward collection latency, indicating degradation nor sex significantly affected motivation during sessions.

In human versions of the CPT, response time variability is often used as a measure of performance alongside sensitivity measures and correlates with clinical severity and drug response in attention-deficit/hyperactivity disorder (ADHD; Levy et al., 2018). Here, we found that stimulus degradation increases reaction time variability (*F*_*4, 48*_ = 6.032, *p* = 0.0005) and there were no significant effects of sex (*F*_*1, 12*_ = 0.0160, *p*= 0.9014) or degradation X sex effects (*F*_*4, 48*_ = 1.575, *p* = 0.1962; Figure 3d) on this measure.

## 4. Discussion

In these studies, we show that stimulus degradation decreases sensitivity (d’), slows reaction times, and increases reaction time variability, all measures of impaired performance, in a mouse touchscreen-based CPT. The changes in performance we observed are qualitatively similar to what has been reported in people tested with degraded stimuli versions of the CPT (Nuechterlein et al., 2015; Nuechterlein et al., 1983). These data provide additional evidence for the translational value of this suite of mouse touchscreen-based CPTs.

As demonstrated previously by multiple research groups, mice readily learn and achieve a high level of proficiency in touchscreen-based CPTs (Caballero-Puntiverio et al., 2020; Caballero-Puntiverio et al., 2019; Hvoslef-Eide et al., 2018). Here, we tested mice on a simplified version of the CPT with parameters corresponding to Stage 3 of training in previous datasets. This modification allowed us to decrease total training time and focus on the effects of stimulus degradation on high-baseline performance. We believe this modified protocol can serve as a platform for studying the neural circuits involved in regulating stimulus discrimination in the CPT and screening of novel therapeutic strategies.

In this study, we tested both male and female C57BL/6J mice. We saw no significant differences in the primary measure of performance (d’) in testing phases, though females did take longer to reach their performance plateau. Additionally, there were no sex differences in the effects of stimulus degradation. Interestingly, we did see significant differences in the response strategy utilized throughout training and testing, independent from stimulus degradation effects. We quantified response strategy using the response criterion measure c from signal detection theory. Females demonstrated a more liberal response strategy throughout the experiment. Liberal responding can readout as a higher hit rate, higher false alarm rate, or some combination of both. In this study, the female mice had significantly higher false alarm rates with no change in hit rates compared to male mice.

The factors underlying the sex difference in response bias in our sample are unclear and human studies in response strategy have shown mixed results. A meta-analysis of children with ADHD and studies of adult participants indicate males have more liberal response biases (Burton et al., 2010; Hasson & Fine, 2012). This difference may be due to a species disconnect or differences in the CPT test parameters. Studies in humans with direct comparisons between males and females have used the Conners’ CPT, which has a very high (>80%) target presentation rate, similar to the Five-Choice Continuous Performance Task (5C-CPT) which is also used with mice (Young, Light, Marston, Sharp, & Geyer, 2009). Due to the high target rate, the Conners’ CPT is thought to be a more sensitive measure of response inhibition where CPTs with lower target rates are thought to be more sensitive measures of vigilance (Ballard, 2001). These differences in task parameters may produce different response strategies and biases. Future studies will need to address the effect of sex on performance in versions of the rCPT that differ on the S+ probability rate and compare those results to human versions with low target probability rates.

Reaction time and reaction time variability, measured as the standard deviation of the reaction time, across a CPT session are sensitive measures of overall performance and increased variability is correlated with impaired performance and treatment response (Fredriksen, Egeland, Haavik, & Fasmer, 2021; Levy, Pipingas, Harris, Farrow, & Silberstein, 2018). To our knowledge, reaction time variability has not been reported in previous studies utilizing the rCPT. Here, we show that both reaction time and reaction time variability increase with the level of stimulus degradation and are correlated with impaired performance. Additional studies are required to determine if reaction time is responsive to treatments that improve performance in mice.

The current studies provide additional evidence that the rCPT is a behavioral assay with significant translational utility. Specifically, mice perform similarly to humans in response to degraded stimuli. Therefore, degraded stimuli versions of the rCPT have the potential to provide greater flexibility for the identification of neurophysiological biomarkers and preclinical screening of novel cognitive enhancers.

## Data Availability Statement

The datasets generated by this project are available upon request.

## Acknowledgements

The authors thank Michael Noback for technical assistance.

## References

Ballard, J. C. (2001). Assessing attention: comparison of response-inhibition and traditional continuous performance tests. J Clin Exp Neuropsychol, 23(3), 331–350. doi: 10.1076/jcen.23.3.331.1188

Beck, L. H., Bransome, E. D., Jr., Mirsky, A. F., Rosvold, H. E., & Sarason, I. (1956). A continuous performance test of brain damage. J Consult Psychol, 20(5), 343–350. doi: 10.1037/h0043220

Berger, I., Slobodin, O., & Cassuto, H. (2017). Usefulness and Validity of Continuous Performance Tests in the Diagnosis of Attention-Deficit Hyperactivity Disorder Children. Arch Clin Neuropsychol, 32(1), 81–93. doi: 10.1093/arclin/acw101

Berman, S. M., Kuczenski, R., McCracken, J. T., & London, E. D. (2009). Potential adverse effects of amphetamine treatment on brain and behavior: a review. Mol Psychiatry, 14(2), 123–142. doi: 10.1038/mp.2008.90

Borgaro, S., Pogge, D. L., DeLuca, V. A., Bilginer, L., Stokes, J., & Harvey, P. D. (2003). Convergence of different versions of the continuous performance test: clinical and scientific implications. J Clin Exp Neuropsychol, 25(2), 283–292. doi: 10.1076/jcen.25.2.283.13646

Burton, L., Pfaff, D., Bolt, N., Hadjikyriacou, D., Silton, N., Kilgallen, C., … Allimant, J. (2010). Effects of gender and personality on the Conners Continuous Performance Test. J Clin Exp Neuropsychol, 32(1), 66–70. doi: 10.1080/13803390902806568

Caballero-Puntiverio, M., Lerdrup, L. S., Arvastson, L., Aznar, S., & Andreasen, J. T. (2020). ADHD medication and the inverted U-shaped curve: A pharmacological study in female mice performing the rodent Continuous Performance Test (rCPT). Prog Neuropsychopharmacol Biol Psychiatry, 99, 109823. doi: 10.1016/j.pnpbp.2019.109823

Caballero-Puntiverio, M., Lerdrup, L. S., Grupe, M., Larsen, C. W., Dietz, A. G., & Andreasen, J. T. (2019). Effect of ADHD medication in male C57BL/6J mice performing the rodent Continuous Performance Test. Psychopharmacology (Berl), 236(6), 1839–1851. doi: 10.1007/s00213-019-5167-x

Fredriksen, M., Egeland, J., Haavik, J., & Fasmer, O. B. (2021). Individual Variability in Reaction Time and Prediction of Clinical Response to Methylphenidate in Adult ADHD: A Prospective Open Label Study Using Conners’ Continuous Performance Test II. J Atten Disord, 25(5), 657–671. doi: 10.1177/1087054719829822

Green, M. F., Kern, R. S., Braff, D. L., & Mintz, J. (2000). Neurocognitive deficits and functional outcome in schizophrenia: are we measuring the “right stuff”? Schizophr Bull, 26(1), 119–136. doi: 10.1093/oxfordjournals.schbul.a033430

Green, D.M., & Swets, J.A. (1966). Signal detection theory and psychophysics. John Wiley.Robyn

Grier, R. A., Warm, J. S., Dember, W. N., Matthews, G., Galinsky, T. L., & Parasuraman, R. (2003). The vigilance decrement reflects limitations in effortful attention, not mindlessness. Hum Factors, 45(3), 349–359. doi: 10.1518/hfes.45.3.349.27253

Hasson, R., & Fine, J. G. (2012). Gender differences among children with ADHD on continuous performance tests: a meta-analytic review. J Atten Disord, 16(3), 190–198. doi: 10.1177/1087054711427398

Huntley, J. D., Hampshire, A., Bor, D., Owen, A. M., & Howard, R. J. (2017). The importance of sustained attention in early Alzheimer’s disease. Int J Geriatr Psychiatry, 32(8), 860–867. doi: 10.1002/gps.4537

Hvoslef-Eide, M., Nilsson, S. R., Hailwood, J. M., Robbins, T. W., Saksida, L. M., Mar, A. C., & Bussey, T. J. (2018). Effects of anterior cingulate cortex lesions on a continuous performance task for mice. Brain Neurosci Adv, 2. doi: 10.1177/2398212818772962

Kim, C. H., Hvoslef-Eide, M., Nilsson, S. R., Johnson, M. R., Herbert, B. R., Robbins, T. W., … Mar, A. C. (2015). The continuous performance test (rCPT) for mice: a novel operant touchscreen test of attentional function. Psychopharmacology (Berl), 232(21-22), 3947–3966. doi: 10.1007/s00213-015-4081-0

Koetsier, G. C., Volkers, A. C., Tulen, J. H., Passchier, J., van den Broek, W. W., & Bruijn, J. A. (2002). CPT performance in major depressive disorder before and after treatment with imipramine or fluvoxamine. J Psychiatr Res, 36(6), 391–397. doi: 10.1016/s0022-3956(02)00026-2

Levy, F., Pipingas, A., Harris, E. V., Farrow, M., & Silberstein, R. B. (2018). Continuous performance task in ADHD: Is reaction time variability a key measure? Neuropsychiatr Dis Treat, 14, 781–786. doi: 10.2147/NDT.S158308

Liu, J., Liu, B., Wang, M., Ju, Y., Dong, Q., Lu, X., … Li, L. (2021). Evidence for Progressive Cognitive Deficits in Patients With Major Depressive Disorder. Front Psychiatry, 12, 627695. doi: 10.3389/fpsyt.2021.627695

Liu, S. K., Chiu, C. H., Chang, C. J., Hwang, T. J., Hwu, H. G., & Chen, W. J. (2002). Deficits in sustained attention in schizophrenia and affective disorders: stable versus state-dependent markers. Am J Psychiatry, 159(6), 975–982. doi: 10.1176/appi.ajp.159.6.975

Nuechterlein, K. H., Edell, W. S., Norris, M., & Dawson, M. E. (1986). Attentional vulnerability indicators, thought disorder, and negative symptoms. Schizophr Bull, 12(3), 408–426. doi: 10.1093/schbul/12.3.408

Nuechterlein, K. H., Green, M. F., Calkins, M. E., Greenwood, T. A., Gur, R. E., Gur, R. C., … Braff, D. L. (2015). Attention/vigilance in schizophrenia: performance results from a large multi-site study of the Consortium on the Genetics of Schizophrenia (COGS). Schizophr Res, 163(1-3), 38–46. doi: 10.1016/j.schres.2015.01.017

Nuechterlein, K. H., Parasuraman, R., & Jiang, Q. (1983). Visual sustained attention: image degradation produces rapid sensitivity decrement over time. Science, 220(4594), 327–329. doi: 10.1126/science.6836276

Riccio, C. A., Reynolds, C. R., Lowe, P., & Moore, J. J. (2002). The continuous performance test: a window on the neural substrates for attention? Arch Clin Neuropsychol, 17(3), 235–272.

Young, J. W., Light, G. A., Marston, H. M., Sharp, R., & Geyer, M. A. (2009). The 5-choice continuous performance test: evidence for a translational test of vigilance for mice. PLoS One, 4(1), e4227. doi: 10.1371/journal.pone.0004227

